# Glucose lowering effects by sago (*Metroxylon sagu Rottb*) resistant starches Type 2 and 4 in spontaneously type 2 diabetes, Goto kakizaki rat

**DOI:** 10.1101/2022.07.26.501535

**Authors:** Ezarul Faradianna Lokman, Sal Hazreen Bugam, Aina Shafiza Ibrahim, Nurleyna Yunus, Fazliana Mansor, Vimala Balasubramaniam, Khairul Mirza Mohamad, Rabizah Md Lazim, Awang Zulfikar Rizal Awang Seruji

## Abstract

The beneficial effects of resistant starch (RS) consumption on health in terms of reducing postprandial hyperglycemia are evident. However, the potential of local Sarawak sago RS in regulating glucose has not been extensively studied. This study aims to identify glucose lowering effects of Sarawak sago RS namely, native unmodified (RS2) and chemically modified (RS4). An oral glucose tolerance test was performed before and after one-month treatment with sago RS2 and RS4 in spontaneously type 2 diabetes, goto kakizaki rat. The mechanisms involved were further explored by screening the in vitro inhibitory activities of α-Glucosidase and DPP-IV. Histopathology examination for pancreas, kidney and liver tissues was performed in response to sago RS treatment using hematoxylin and eosin (H&E) staining.The blood glucose area under the curve (AUC) in RS-treated groups was decreased and significant in RS2-treated group (p<0.05). Improved insulin AUC and higher glucagon like peptide (GLP-1) levels were observed in all RS-treated groups (p<0.05). Sago RS2 and RS4 might have potential roles in regulating glucose via α-Glucosidase and DPP-IV inhibitory activities by reducing glucose absorption in the intestine. For histopathology study, although insignificant, sago RS2 and RS4 attenuated lesion scores of pancreatic tissue whereas the liver and kidney tissues significantly showed lesser lesion scores as compared to the control diabetic group suggesting the potential of RS in reducing cell degeneration which indeed requires further validation.Findings of this study suggests the therapeutic potential of sago RS in the T2D management which may justify further investigation to be done.

## Introduction

Type 2 diabetes (T2D) is characterized by hyperglycemia associated with diminished insulin secretion and insulin resistance which leads to increased hepatic glucose output and decreased glucose peripheral utilization (1). Uncontrolled hyperglycemia results in several complications such as macrovascular (coronary artery disease, peripheral arterial disease and stroke) and microvascular (neuropathy, nephropathy and retinopathy) (2). Postprandial hyperglycemia is influenced by several factors including gastric emptying process, intestinal glucose absorption rate, insulin sensitivity in peripheral tissues, glycogenolysis process after meals, hepatic gluconeogenesis and insulin secretion rate during postprandial period. Postprandial hyperglycaemia is defined as a plasma glucose level more than 7.8 mmol/L (140 mg/dL) 1-2 hours after food ingestion (3). Current oral treatment for type-2 diabetes are targeted at reducing hepatic glucose output, improving insulin release, improving glucose absorption and increasing peripheral glucose utilization (4). Physical activity along with a dietary regime have been suggested to be important in regulating blood glucose levels, thus important for T2D management (5,6).

In terms of dietary intake, resistant starch (RS) consumption with low glycaemic index properties has been shown to be beneficial in managing postprandial hyperglycemia. The process whereby RS skips digestion in the small intestinal and is fermented by microbial in the colon preventing glucose spike from occurring (7). In principle, the more the RS content, the slower the digestion rate and the lower is the glycaemic index (GI) which determines the ability of food to raise the blood glucose level (8). The postprandial glucose excursion in people with diabetes can be elevated and prolonged, therefore consuming resistant starch may help control glucose level. The glycaemic effect of food depends on numerous factors affecting the formation of RS such as the structure of amylose/ amylopectin ratio, processing factors, formation in processed food, the native environment of the starch granule and others Thus, the factors affecting the GI values depends on the RS formation (9,10).

Dipeptidyl peptidase (DPP)-IV and α-glucosidase inhibitors are two classes of oral drugs used for the hyperglycaemic treatment in type 2 diabetes (11) by decreasing the postprandial hyperglycemia via digestive system (12). α-Glucosidase inhibitor which is involved the final step in the digestive process of carbohydrates suppresses postprandial hyperglycemia by retarding the glucose release from dietary complex carbohydrates (13) and prevents the digestion of carbohydrates (12). Inhibition of α-glucosidase in the intestine causes the spread of carbohydrate digestion process to the lower part of small intestine which delays the overall glucose absorption rate into the blood which in turn decrease the postprandial rise in blood glucose (13).

Certain food-derived peptides promote glucose homeostasis by regulating sugar absorption and insulin levels through inhibition of dipeptidyl peptidase IV (DPP-IV). DPP-IV is a brush-membrane-associated prolyl dipeptidyl peptidase that is involved in the in vivo hydrolysis of incretins. DPP-IV is the enzyme responsible for the proteolytic breakdown of incretins, which play an important role in the regulation of glucose homeostasis (14). Incretins (intestinal insulinotropic hormones) are primarily responsible for regulating glucose levels. Two peptide incretin hormones involved in blood glucose control have been identified in humans, namely glucose-insulinotropic peptide (GIP) and glucagon-like peptide-1 (GLP-1). They are released from the gut in response to food intake and exert a strong insulinotropic effect (contribute to lowering glucose levels by stimulating insulin secretion and inhibiting glucagon release), helping to control postprandial glucose levels (14,15). So far, the presence of DPP-IV inhibitory (DPP-IV) peptides in food products of animal origin has been demonstrated [10].

Several studies reported significant changes in postprandial glucose and the insulin incremental area under the curve (iAUC) after consuming RS in prediabetes and diabetes adults (16–20). For *in vivo* studies, the glucose-lowering effects by different sources of RS were evident in diabetic (alloxn/streptozotcin-diabetic induced, Goto kakizaki, Zucker diabetic fatty (ZDF) rats) and healthy rodents (Wistar rats, Sprague Dawley) (21–29). In terms of pancreatic islets study, Shen et al., 2011 found a significant increase of β-cell density in diabetic goto kakizaki rats fed with RS (29). The glucose regulating effect of several types of RS with Dipeptidyl peptidase (DPP-IV) and α-glucosidase inhibitory activities was previously demonstrated in both healthy and disease models of several studies (12,30– 33). Furthermore, using Rat Glucose Metabolism RT^2^ Profiler PCR Array, our previous published data have shown several genes associated with glucose and glycogen metabolism pathways were up- and downregulated in the liver of diabetic rats treated with sago RS2 and RS4 (34). Despite many promising results obtained from several studies on RS, the beneficial effects of our local RS extracted from sago (*Metroxylon sagu Rottb*.) in terms of managing blood glucose are still limited and lacking. This justifies the extension of studying the potential of our local Sarawak sago RS to be carried out in human with T2D.

In Malaysia, Sarawak produces the world’s biggest sago since in the 1970s’ (35,36) which can be found primarily at nearby river areas Mukah and Betong divisions. The amylose/amylopectin ratio of Sago are reported to be higher (24 – 31%) as compared to short-medium grain rice (15 – 18%) and glutinous rice (4%) which are commonly consumed in Asia (36,37). RS content of sago native starch was reported to contain up to 68.99% (38). Chemical modification such as phosphorylation, acetylation, hydroxypropylation is known to increase the RS content of a starch (39). Acetylation has shown to increase the RS content of sago native starch to up to 74% - which is comparable to the commercially available wheat RS from Fibersym™ (38).

In Sarawak, various types of food have been produced from sago and widely used in food industries including sago pearls, tabaloi (a traditional delicacy biscuit), keropok (shrimp crackers), puddings, jellies and used as thickener in food products and made into noodles and vermicelli (36). Starch in food may be classified to three categories namely rapidly digestible starches, slowly digestible starches, and resistant starches based on the results of *in vitro* digestion (40). RS can be further classified according to the reason for resistance to digestion. There are 5 types of RS known as RS1 (Physically inaccessible starch-coarsely ground or whole-kernel grains), RS2 (Granular starch with the B- and C-polymorph-high amylose maize starch, raw potato, raw banana starch), RS3 (Retrograded starch-cooked and cooled starchy foods), RS4 (Chemically modified starches-cross-linked starch and octenyl succinate starch) and RS5 (Amylose-lipid complex-stearic acid-complexed high-amylose starch) (9).

This current study was carried out to investigate the postprandial glucose lowering effects of local Malaysian sago native (RS2) and modified (RS4) from Sarawak. The mechanisms involved associated with inhibitory activities of α-Glucosidase and dipeptidyl peptidase IV (DPP-IV) in response to the Sarawak sago RS were accessed, and thus may identify the importance of RS in managing postprandial hyperglycaemia.

## Materials and methods

### Analysis of Sago Starches

Sago native starch (Sago RS2) used in the study was obtained from local supplier. A portion of this native starch was then subjected to further chemical modification to produce sago RS4 which was obtained from a separate project. The sago modified starch used in this study was a cross-linked dual modified sago starch, which was obtained by treating the native sago starch with sodium trimethaphosphate (STMP) and sodium tripolyphosphate (STPP). This modification has resulted in a type 4 resistant starch (RS4), with the degree of substitution of 0.129. The RS content has also increased considerably from its initial native starch RS of 45.53, to 60.66 after this modification. This starch modification was aimed at increasing the RS content of the native starch. The resistant corn starch is a commercial HiMaize RS2 starch (LifeSource resistant corn Starch 260), while the regular corn starch was sourced from local grocery store. The assay for the RS content for these starches was conducted using Megazym resistant starch assay kit (Megazym, UK) adopting AOAC methods 2002.02. The RS percentage for Sarawak sago native starch (Sago RS2), Sarawak sago modified starch (Sago RS4) and Resistant corn starch 260 (HM) are 45.53 ± 0.24, 60.66 ± 0.39 and 41.64 ± 0.89 respectively. The Scanning electron microscope (SEM) images of sago cross-linked dual modified starch granule (RS4) and sago native starch granule (RS2) are shown in **Figure 1a** and **Figure 1b** respectively.

**Figure 1:**
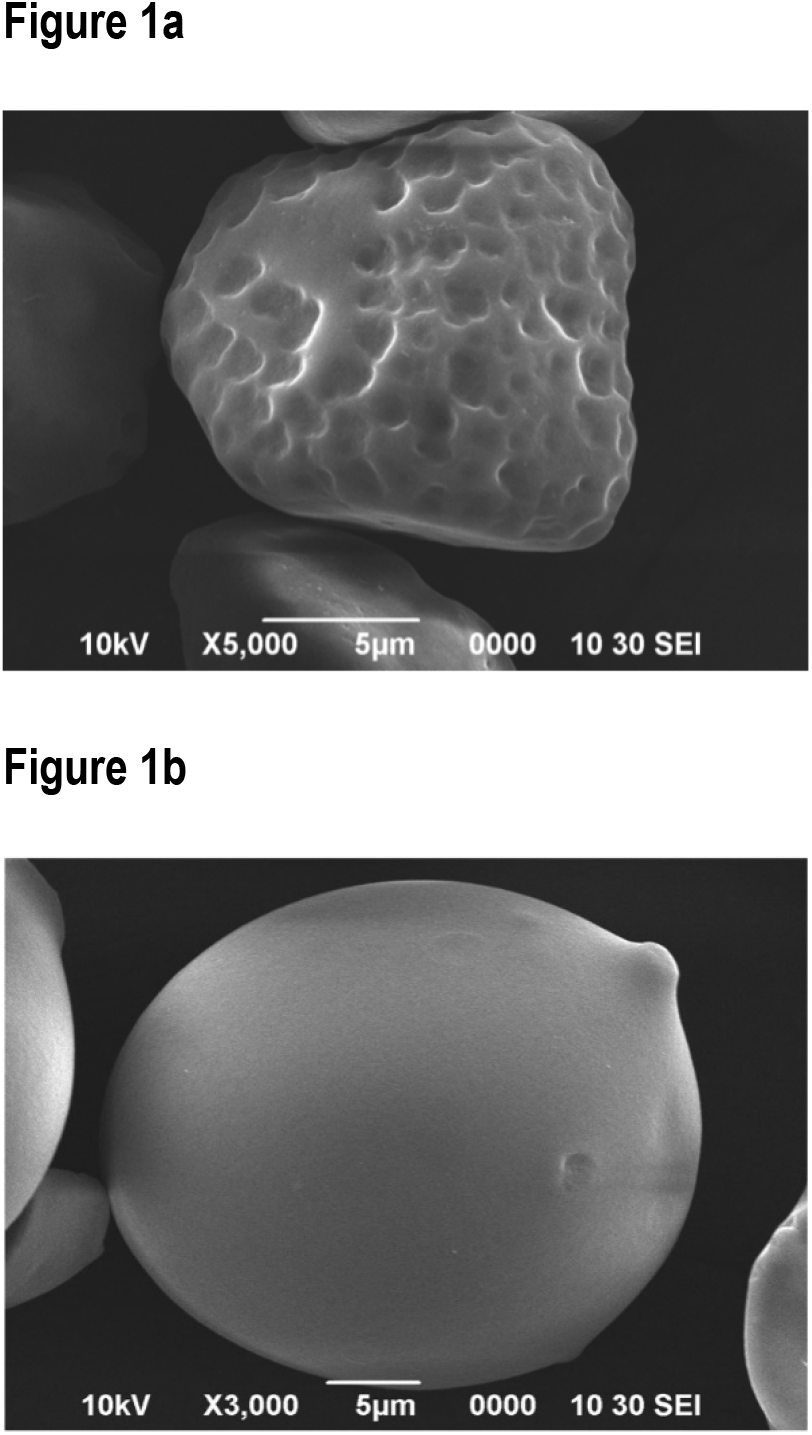
Scanning electron microscope (SEM) images of (a) sago cross-linked dual modified starch granule (RS4) observed at x5,000 magnification and (b) sago native starch granule (RS2) observed at x3,000 magnification.

### *In vivo*: one month treatment with Sago Dietary Resistant Starch and oral glucose tolerance test (OGTT) in Goto kakizaki rats

Healthy Goto kakizaki (GK) male rats aged 8 weeks with mean body weight 125.8±8.8 g were purchased from Clea Japan, Inc. The GK rat is a lean model of type 2 diabetes which develop mild hyperglycaemia early in life due to defect in β -cell mass (41). The animals were quarantined for 2 weeks and acclimatized for 5 days prior to start of the study. The animals were placed in individual ventilated cages, supplied with reverse osmosis drinking water (Sartorius, Germany) and a commercial rat pellet (Specialty Feeds, Australia), ad libitum with controlled temperature at (20± 2°C), 40-60% humidity under 12 hours of light and dark cycle. Ethical approval for this study was obtained from the Animal Care and Use Committee, Ministry of Health Malaysia (ACUC/KKM/02(03/2018) and all experiments were conducted in accordance to the regulations.

The rats reached 11 weeks during the start of the experiment with mean body weight 237.6±13.2 g. The rats were randomly assigned to four groups in individual cages (n=8). Group 1: Control diabetic (CD) (Vehicle); Group 2: Positive control (Hi Maize)(0.07g/kg of b.w); Group 3: Treatment (Sago RS2)(0.4g/kg of b.w); Group 4: (Sago RS4) (0.07g/kg of b.w). Several parameters such general behaviour, body weight changes, food and water consumption were recorded throughout the study.

Glucose tolerance test was performed before and after treatment to identify how efficient cells can process sugar. Rats were left fasted overnight prior to the test. Fasting blood glucose level for all rats were measured prior to glucose load (0.2g/kg of body weight) by tail prick using a glucometer (Accucheck Aviva Plus, Roche, USA). Blood glucose level was measured at 0, 30, 60 and 120 min using a glucometer (Accucheck Aviva Plus, Roche, USA). Blood was also collected at 0, 30, 120 min in heparin tubes for insulin content from plasma measurement. Blood was then centrifuged at 2500 x g for 10 min and plasma was transferred into 1.5 ml Eppendorf tubes and kept at -20°C. After a month treatment, rats were anesthetized using isofluorene supplied with oxygen and blood was withdrawn by cardiac puncture method followed by immediate euthanize. Blood was collected in EDTA tube, centrifuged at 2000 x g for 10 min and plasma was transferred into 1.5 ml Eppendorf tubes and kept at -20°C for. Plasma was used for analyte analysis (Ghrelin, glucagon like peptide (GLP-1) and glucagon) using ELISA kit (Elabscience, China).

### *In vitro*: Effects of Sago Resistant Starch on inhibitory activities

#### DPP-IV inhibition Assay

DPP-IV activity was analysed using a DPP-IV inhibitor screening assay kit (Cayman Chemical, USA) which provided a fluorescence-based method for screening DPP-IV inhibitor. This assay was performed based on the standard protocol with slightly modification by using the 96 well microplates. A pre-incubation volume reagent prepared contained 30 ml of 20 mM Tris-HCL buffer pH 8.0, 600µl of DPP-IV (human recombinant) enzyme, 3ml of DPP substate, 500µl of sitagliptin positive control inhibitor and various concentration of test material/references inhibitor.

Briefly, the test samples (10 µl) were pipetted onto 96 well microplate (Cayman Chemical, USA) containing fluorogenic substrate, Gly-Pro-Aminomethylcoumarin, DPP substrate (50 µl, final concentration 100 µM). The negative control contained 20mM Tris-HCL buffer pH 8.0 (50 µl) and the reaction substrate Gly-Pro-Aminomethylcoumarin. The reaction was initiated by the addition of DPP -IV (10 µl). Sitagliptin was used as a positive control. Sago RS2 and RS4 have been tested at different concentrations (0.1, 1 and 10 mg/ml). Sitagliptin is a dipeptidyl peptidase-4 inhibitor which control blood glucose level in type 2 diabetes mellitus patients (42). This mixture was incubated at 37^0^C for 30 minutes. The experiment was performed in triplicate and the absorbance in fluorescence was measured at excitation wavelength 355 nm and emission wavelength 460 nm using an BMG LABTECH 96 plate reader (Omega series). DPP-IV IC_50_ values were determined by plotting the percentages of inhibition as a function of the concentration of test compound. The percentage inhibition was calculated using the formula;

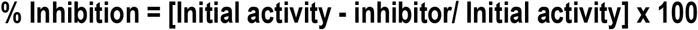

### α-Glucosidase inhibition Assay

α-Glucosidase inhibitory activity was carried out according to the standard method with minor modification. Sago native, sago modified, hi maize have been tested at different concentrations (0.1, 1 and 10 mg/ml). In a 96 well plate, the reaction mixture containing 80 µl α-Glucosidase assay buffer (BioVision, USA), 10µl α-Glucosidase (BioVision, USA), and 10 µl of varying concentrations of samples were preincubates at room temperature for 15-20 minutes. α-Glucosidase activity was determined by measuring the absorbance at 410nm in kinetic mode for 60 minutes at room temperature using microplate reader (Bio-Tek, USA). Acarbose (BioVision, USA) was used as a positive control of α-Glucosidase inhibitor. Acarbose is used as an oral glucose-lowering drug for treatment or prevention of type 2 diabetes mellitus (43). Each experiment was performed in triplicates. The results were expressed as relative enzyme activity percentage, which was calculated using the formula;

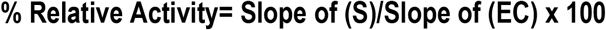

### Histopathology examination for pancreas, kidney and liver

A total of 36 slides consisted of 12 slides each of pancreas, liver and kidneys were presented for histopathology examination at Faculty of Veterinary Medicine, University Putra Malaysia Serdang. The samples were divided into 4 groups, which were groups Control diabetic (CD), Hi Maize (HM), sago RS2 and RS4. Each group consisted of 3 rats and each rat was presented with a section of pancreas and 2 sections each of liver and kidney. All organ sections were stained with haematoxylin and eosin (H&E). Examinations of the pancreas sections were focused on the Islet of Langerhans, particularly the number and size of the Islets and the appearance of the cells. The liver sections were examined for the inflammatory, circulatory and hepatocyte changes while the kidneys were aimed at the changes in glomerulus and convoluted tubules. The identified changes or lesions were described before the extend of the lesions were evaluated and scored as follows:

### Pancreatic lesions

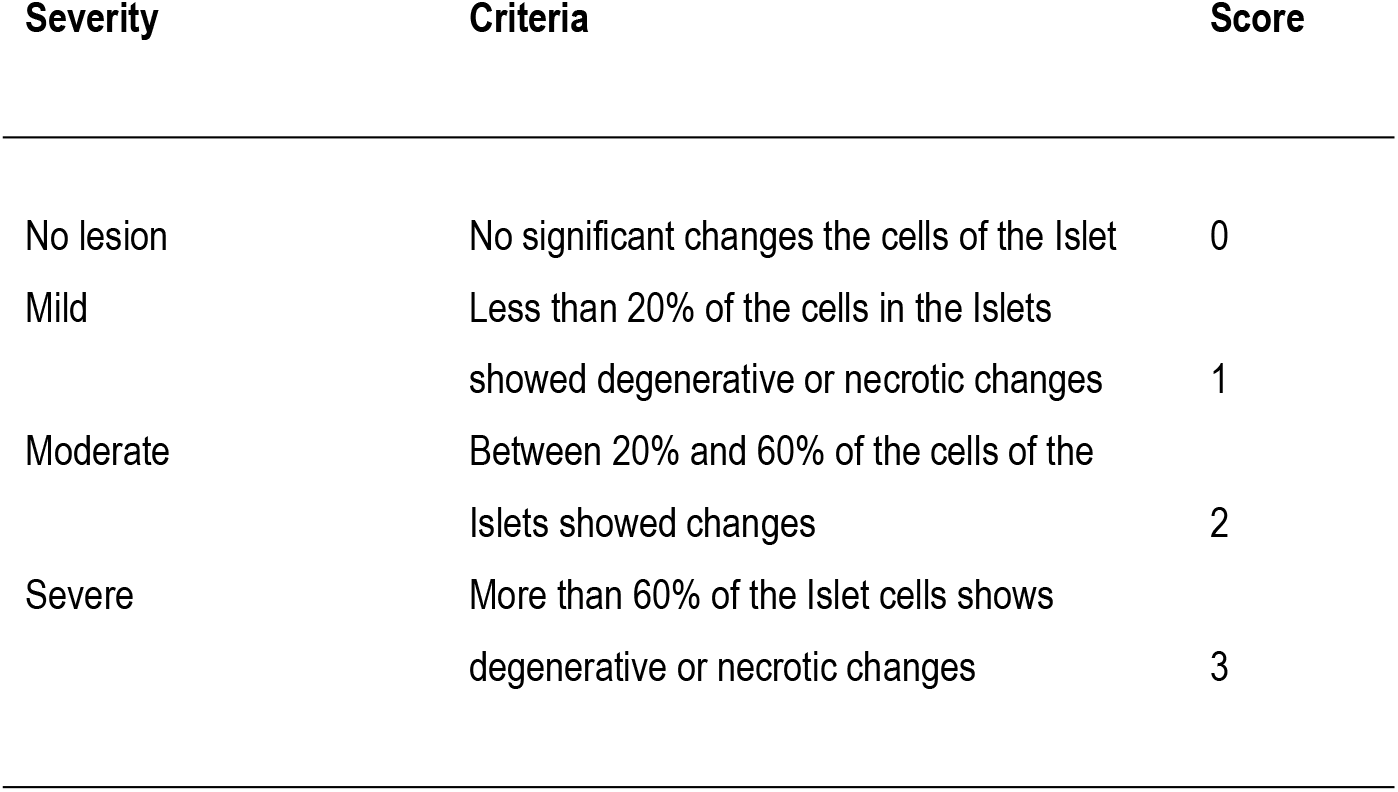

### Liver and kidney lesions

| Severity | Criteria | Score |
| --- | --- | --- |
| No lesion | Absence of the lesion that is being described | 0 |
| Mild lesion | Presence of the lesion involving between 1% and 20% of the organ | 1 |
| Moderate lesion | Presence of the lesion involving between 21% and 60% of the organ | 2 |
| Severe lesion | Presence of the lesion involving more than 60% of the organ | 3 |

## Results

### *In vivo*: one month treatment with Sago Dietary Resistant Starch using Goto kakizaki rats and oral glucose tolerance test (OGTT)

For mortality and morbidity clinical observations, rats were monitored closely during and after treatment for any clinical toxicity signs such as piloerection, diarrhoea, alteration in locomotion, dull fur, emaciation, sedation, soft stool, urination or death. No toxicity signs were observed throughout the study. Before treatment, no significant changes in blood glucose level were observed between different groups. However, after 1 month treatment, blood glucose levels were significantly reduced as compared to CD at min 30 for sago RS2 (p<0.01), min 60 for HM (p<0.05) and sago RS4 (p<0.001) and min 120 for HM (p<0.05); sago RS2 and RS4 (p<0.01) **(Figure 2a)**. The efficacy of treatment based on before and after intervention was also conducted. The blood glucose area under the curve (AUC) following OGTT in the CD group increased significantly on day 28. The rats receiving sago RS2 treatment showed a significant decrease in their blood glucose AUC compared to before they have started of their treatment (p<0.05). No significant difference was reported for sago RS4 and HM however a lowering trend in the rats’ blood glucose AUC were clearly evident for the rats receiving the RS **(Figure 2b)**. AUC for insulin was significant increased by 3.7, 1.2 and 2-fold in groups treated with HM, sago RS2 and RS4 respectively (p<0.05) **(Figure 2c). Table 1** shows several parameters recorded during 1 month treatment such as feed intake, water consumption and body weight changes. No significant changes were observed between groups for feed and water consumption after treatment. Contrary to CD group, sago RS2 consumed the least feed. Treatment with sago RS2 significantly showed the least body weight change (p<0.01) followed by sago RS4 (p<0.05), HM and control.

**Table 1:**
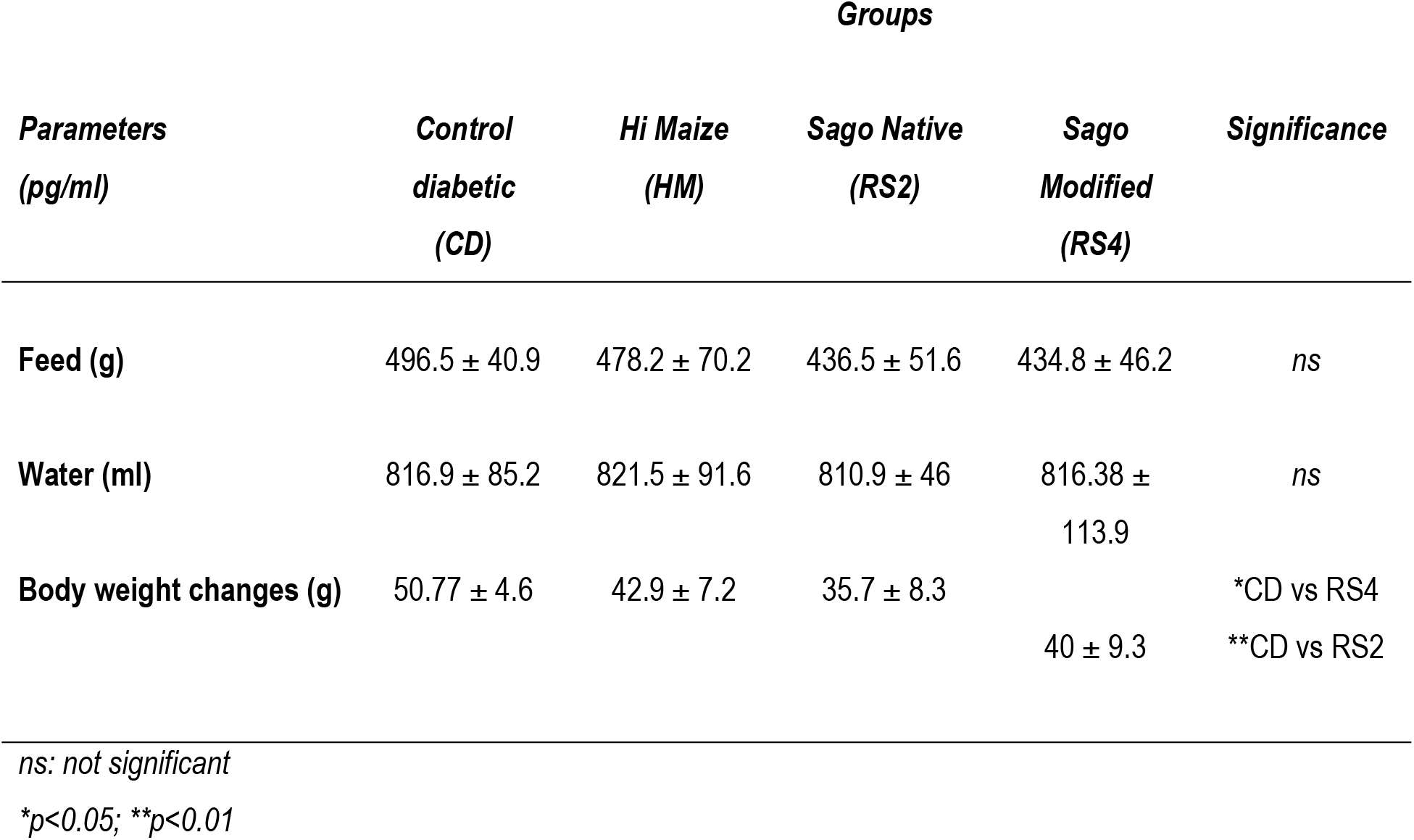
Parameters recorded after 1 month treatment in Goto kakizaki rat.

**Figure 2:**
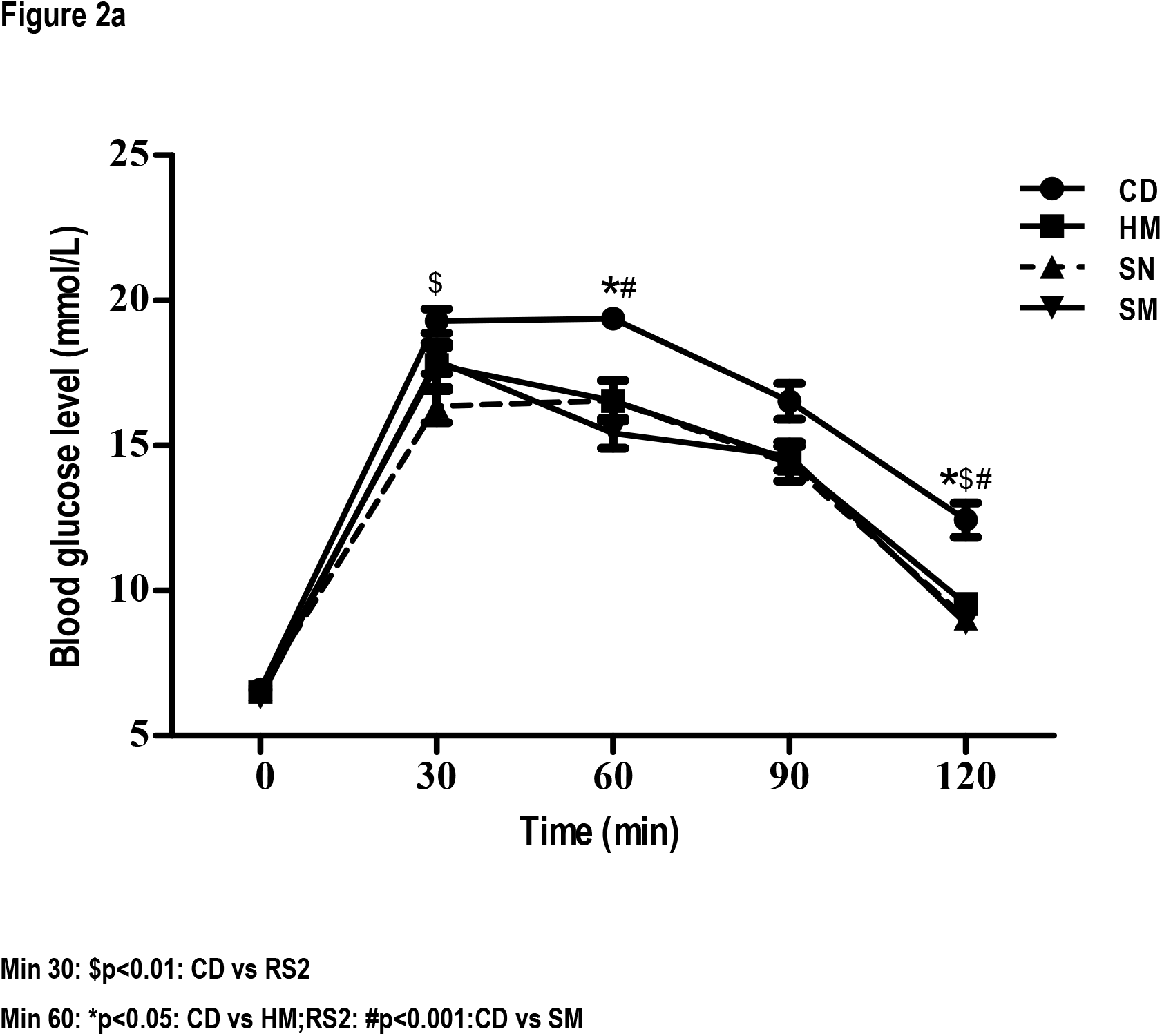
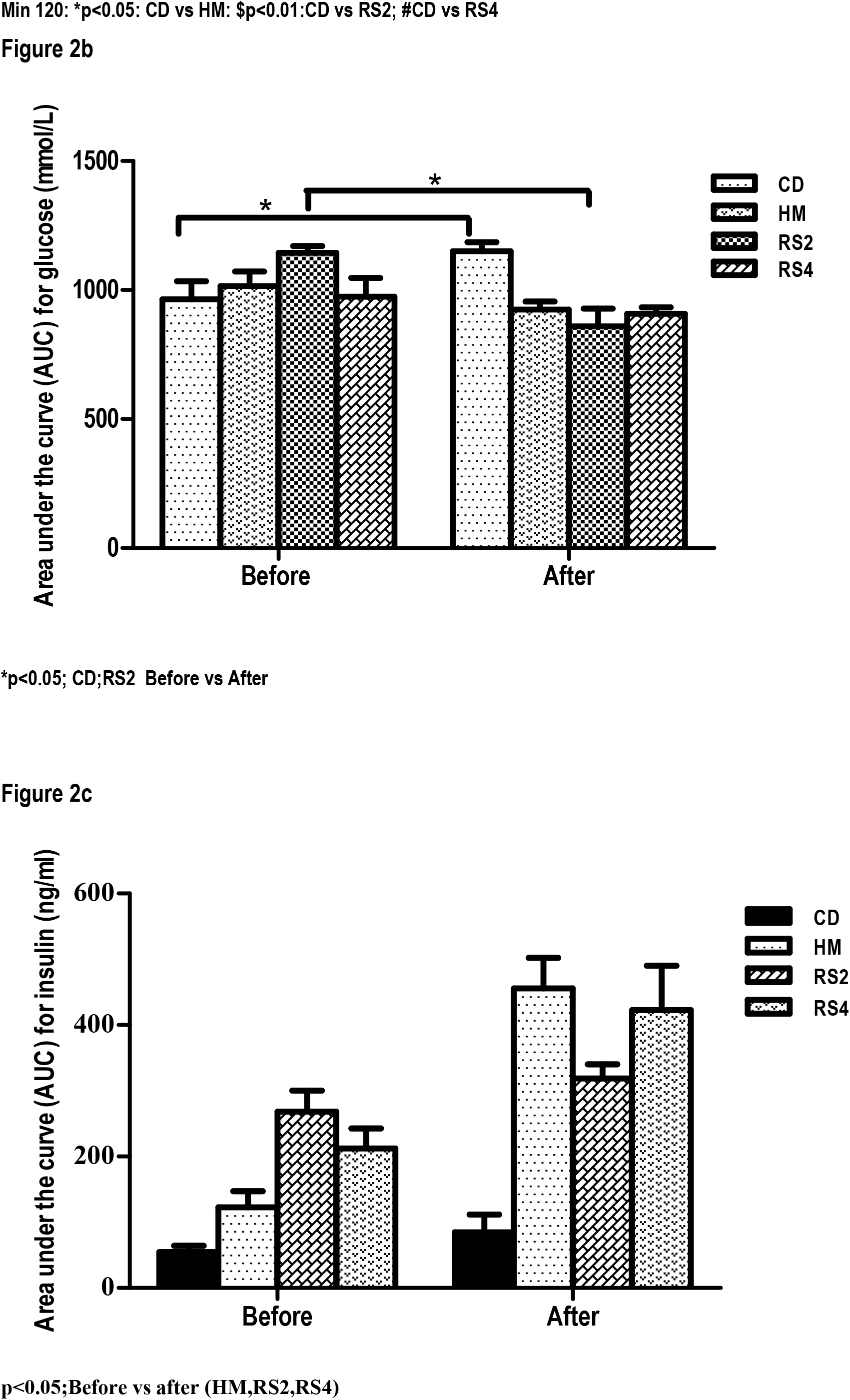
**(a):** Blood glucose levels during oral glucose tolerance test (OGTT) after 1 month treatment in Goto kakizaki rat **(b):** Area under glucose curve (AUC) before and after 1 month treatment in Goto kakizaki rat. **(c):** AUC for insulin after 1 month treatment in Goto kakizaki rat.

**Table 2 shows** several hormones measured using ELISA. **Glucagon Like Peptide-1 (GLP-1):** GLP-1 level was significantly high in rats treated with sago RS2 as compared to control group (p<0.05), followed by HM and sago RS4. GLP-1 is a hormone secreted in the small intestine which increases insulin release and inhibits glucagon secretion, thereby lowering glucose level (44). **Glucagon:** No significant differences was observed for glucagon in all groups despite high level was shown in the sago RS2 group. Glucagon is a peptide hormone, produced by alpha cells of the pancreas to raise the concentration of glucose **Ghrelin levels:** Rats treated with sago RS4 showed the highest Ghrelin level as compared to control. Ghrelin is a hormone produced in the gut to stimulate appetite.

**Table 2:** Hormone levels after 1 month treatment in Goto kakizaki rat.

| **Hormones**<br>**(pg/ml)** | **Groups** |  |  |  | **Significance** |
| --- | --- | --- | --- | --- | --- |
|  | **Control diabetic**<br>**(CD)** | **Hi Maize**<br>**(HM)** | **Sago Native**<br>**(RS2)** | **Sago Modified**<br>**(RS4)** |  |
| **Glucagon Like**<br>**Peptide -1**<br>**(GLP1)** | 625.8 ± 65.6 | 771.5 ± 161.9 | 879.6 ± 117 | 721.35 ± 164.1 | *CD vs RS2 |
| **Glucagon** | 36 ± 2.5 | 37.2 ± 4.4 | 44.7 ± 14.8 | 40.3 ± 5.3 | ns |
| **Ghrelin** | 15.1 ± 14 | 10.3 ± 7.2 | 9.4 ± 8.8 | 25.6 ± 16.7 | ns |

ns: not significant
\*p<0.05

### DPP-IV inhibition Assay

**Figure 3.** showed the inhibitory effect of sago RS2 and RS4 on dipeptidyl peptidase IV (DPP-IV). Dipeptidyl peptidase-4 (DPP-IV) inhibitor play an important role to improve glycaemic control (4). The results showed that RS2 at 1 and 10 mg/ml inhibited DPP-IV activity by 87.72 ± 9.7 and 82.27 ± 5.1%. RS4 only showed DPP-IV inhibitory activity at 10 mg/ml by 87.07 ± 8.9%, whereas HM did not show any inhibitory activity. DPP-IV inhibitory activity by Sitagliptin at 100µM was 87.58 ± 11.16 %. The IC_50_ values of sago RS2 and RS4 against DPP-IV were 0.17 and 5.9 mg/mL, respectively **(Table 3)**.

### α-Glucosidase inhibition assay

The *in vitro* α-glucosidase enzyme activity of sago RS2 and RS4 is shown in **Figure 4**. α-Glucosidase reduces postprandial hyperglycaemia by delaying the process of carbohydrates hydrolysis and glucose absorption (45). All RS displayed inhibitory in a dose dependent manner. At 10 mg/ml, RS2, RS4 and HM exhibited highest α-glucosidase inhibitory activity at 93, 95 and 97% respectively. Acarbose showed complete inhibition against α-glucosidase enzyme. Sago RS4 exhibited the lowest IC_50_ which was 0.6 mg/ml compared to RS2 and HM while IC_50_ of Acarbose was calculated as 0.48 mg/ml **(Table 3)**.

**Table 3:** The IC_50_ values (mg/ml) of Sago RS2, RS4 and Hi Maize for DPP-IV and α-Glucosidase inhibitors

| Sample | IC<sub>50</sub> (mg/ml) |  |
| --- | --- | --- |
|  | DPP-IV | α-Glucosidase |
| Sago RS2 | 0.17±0.0 | 1.4±0.4 |
| Sago RS4 | 5.9±1.4 | 0.6±0.3 |
| Hi Maize (HM) | - | 1.2±0.3 |

**Figure 3:**
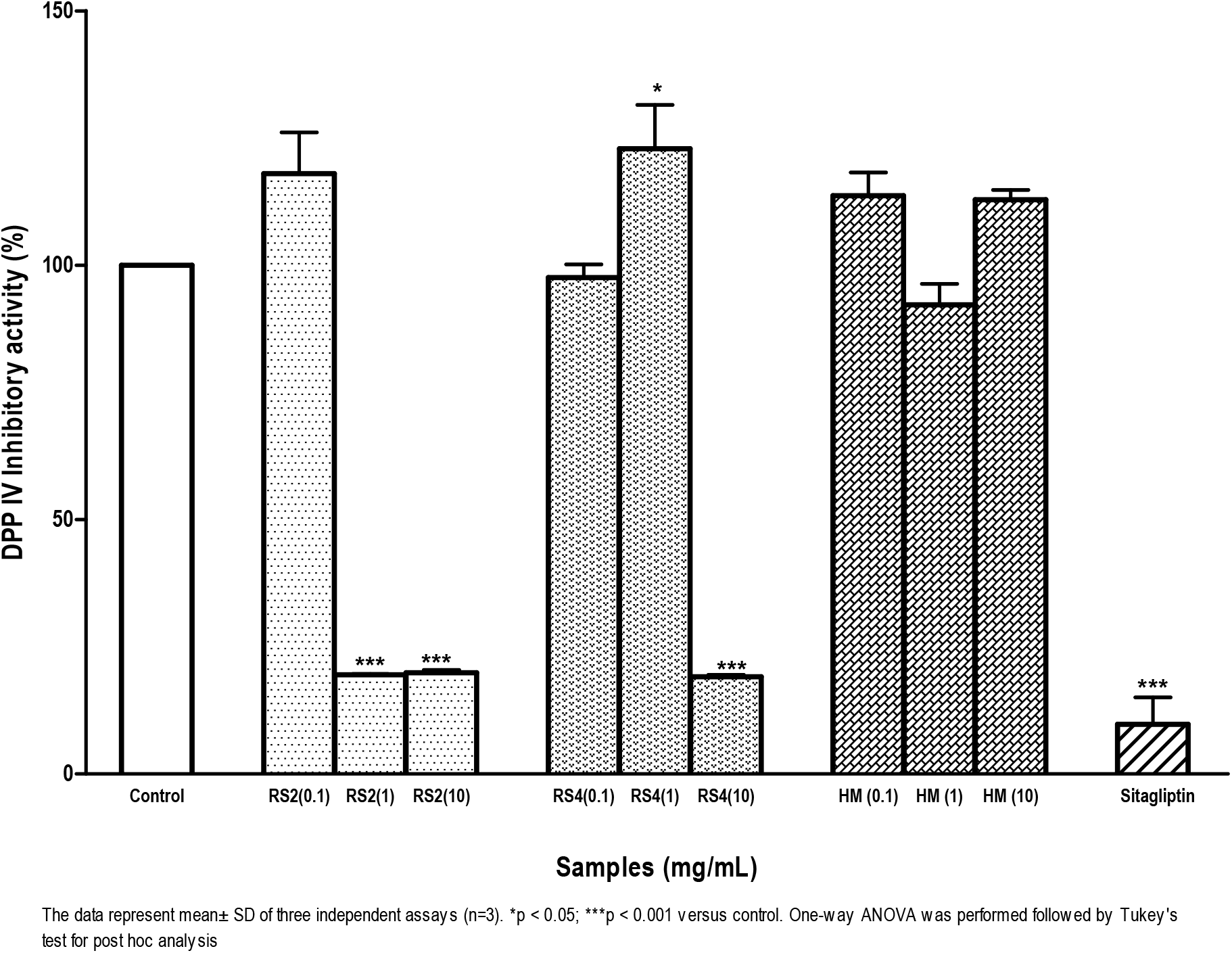
DPP-IV inhibitor activity of sago RS2, RS4 and HM at 0.1, 1, 10 mg/ml concentrations. DPP-IV inhibitor as a control was set at 100 %, and all the other values were normalised to this DPP-IV inhibitor control value, respectively. Sitagliptin was used as a positive control.

**Figure 4:**
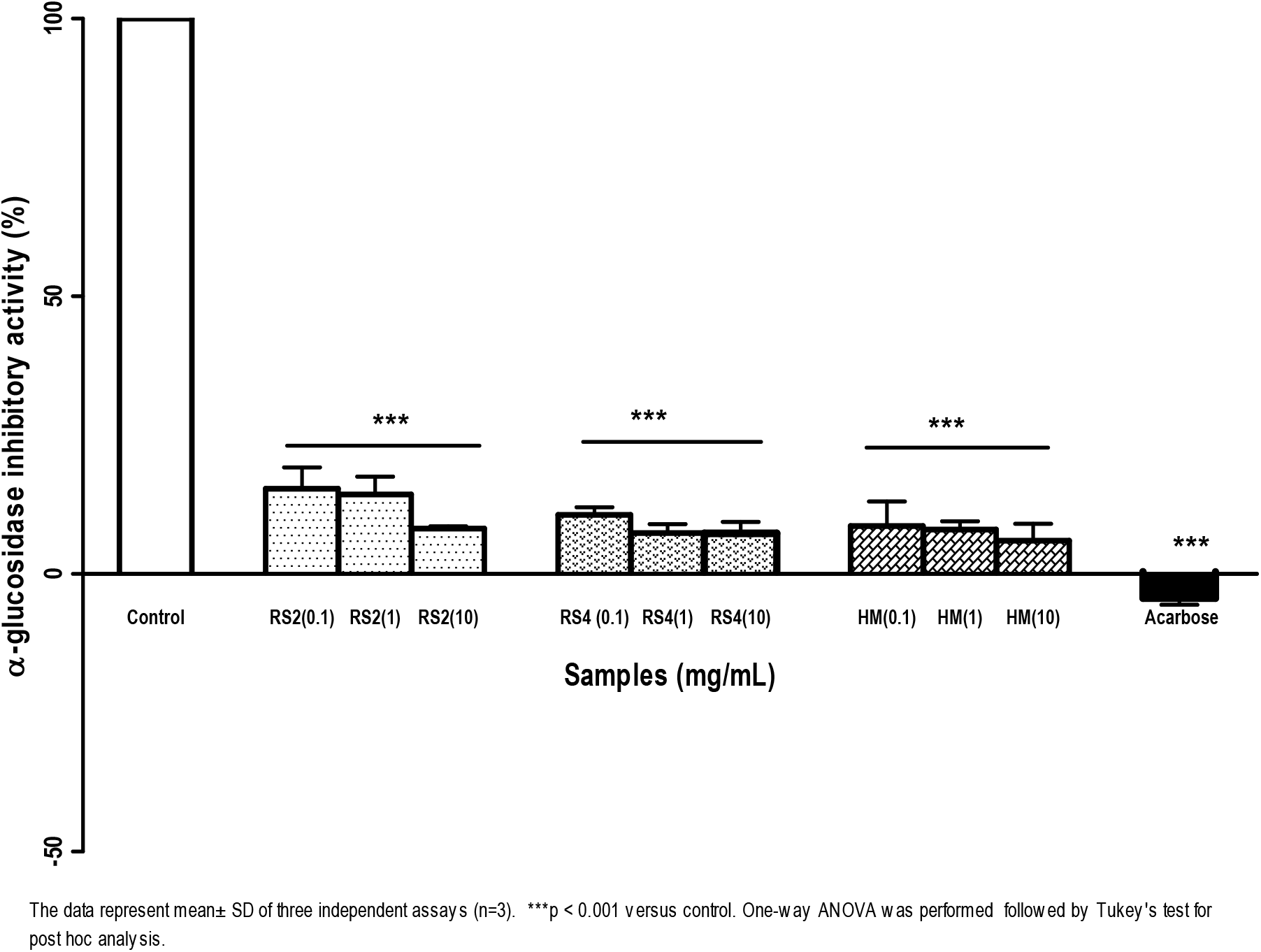
α-Glucosidase inhibitory activity of sago RS2, RS4 and HM at 0.1, 1, 10 mg/ml concentrations. α-Glucosidase inhibitor as a control was set at 100 %, and all the other values were normalised to this α-glucosidase inhibitor control value, respectively. Acarbose was used as a positive control. The data represent mean± SD of three independent assays (n=3). ***p < 0.001 versus control. One-way ANOVA was performed followed by Tukey’s test for post hoc analysis.

### Histopathology examination description and severity of lesions

#### The pancreas

The pancreas of CD group showed moderately congested blood vessels, occasionally with thrombus. The number of Islet of Langerhans ranged between 10 and 14 (average 12) and almost 70% of those were small-sized **(Figure 5a)**. Many cells (about 50%) in the islets were showing pyknotic nuclei, particularly those located in small Islets **(Figure 5b)**. Similarly, the pancreas of group HM showed moderately congested minor blood vessels, occasionally with thrombus. The number Islet of Langerhans ranged between 9 and 11 (average 10), but most were small-sized. Between 50%-60% of the cells in the Islets showed pyknotic nuclei. The blood vessels of group sago RS2 were fairly normal but the number of Islet of Langerhans ranged between 9 and 18 (average 14) and many of those were large-sized. An average of 20% of the cells were pyknotic. Sago RS4 showed relatively mild congestion of the blood vessels, occasionally with thrombus. The number of Islet of Langerhans ranged between 9 and 16 (average 12) and only a few were large-sized. However, the cells of the Islets were fairly normal with occasional few showing pyknosis. All rats showed normal exocrine part of the pancreas. In general, all lesions in the pancreas of all groups showed insignificant (p>0.05) differences, except the extent of islet cellular necrosis of group sago RS4, which is significantly (p<0.05) less extensive than the other groups **(Table 4)**. Nevertheless, cellular necrosis was markedly less severe in group sago RS2 compared to CD group although the difference was insignificant (p>0.05). Although the number and size of the islet of Langerhans differ between groups, they did not significantly (p>0.05) affect the results.

**Table 4:**
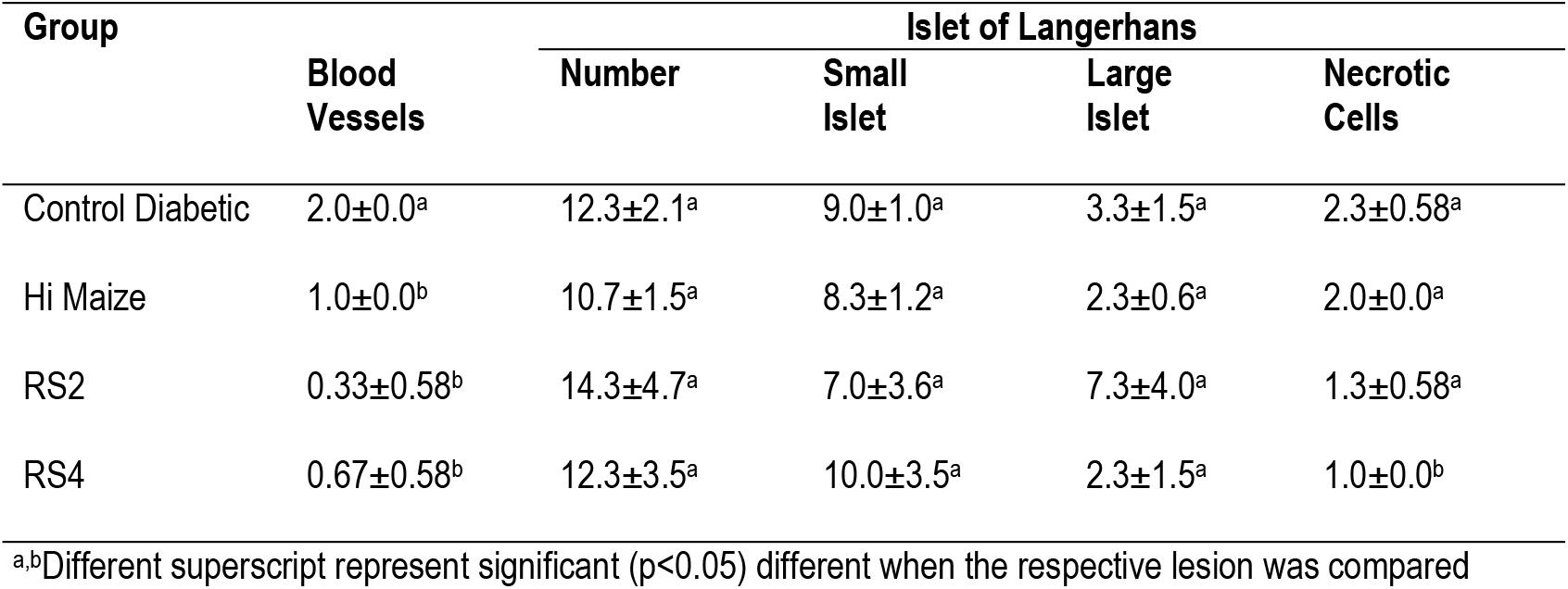
Lesion scores that indicate the severity of lesion in pancreas

**Figure 5(a):**
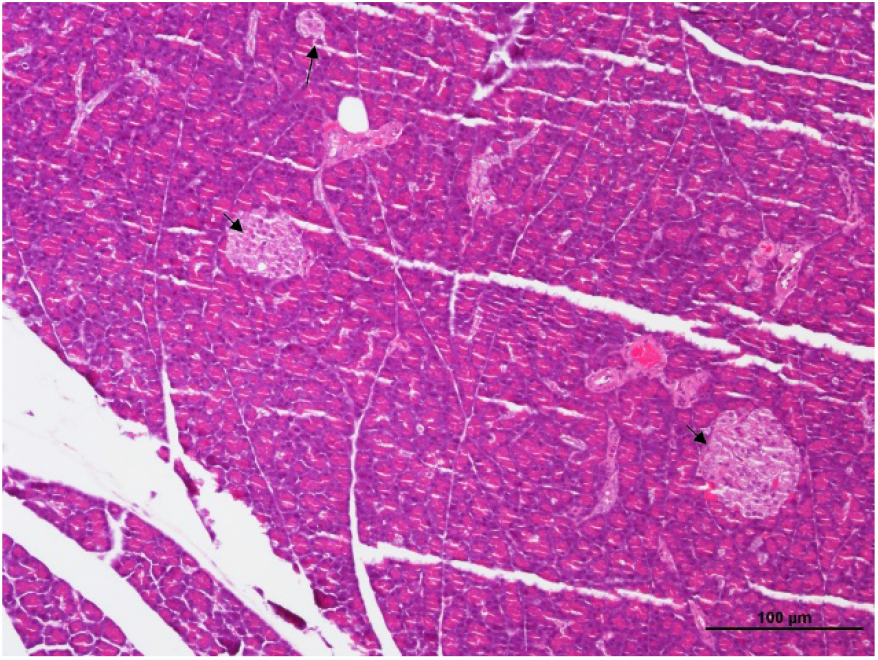
Section of a pancreas from rat of group CD showing two large (arrowheads) and a small islet. HE x40

**Figure 5(b):**
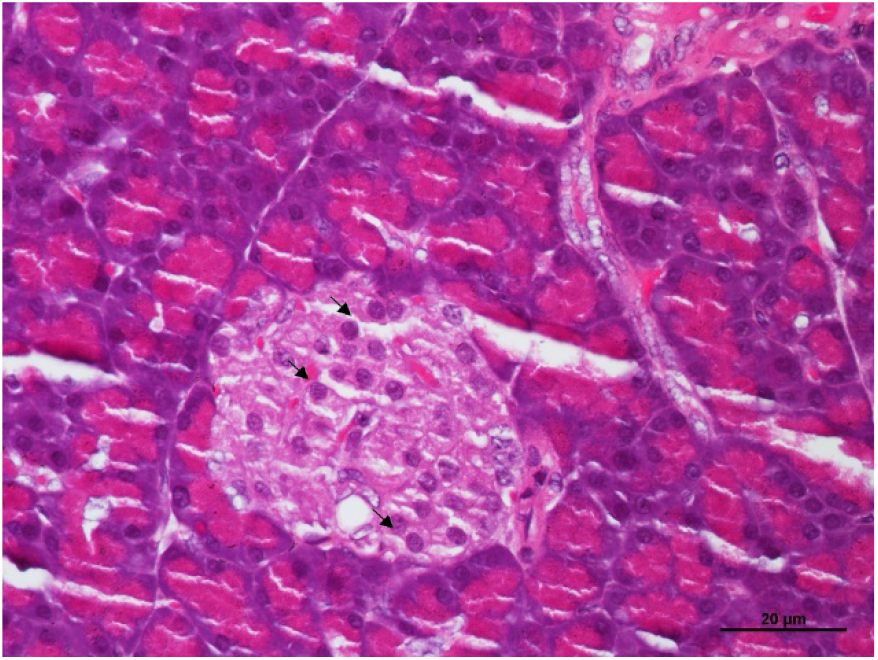
Section of a pancreas from rat of group CD showing cells of the islet. Approximately 50% of the cells showed pyknotic nuclei (arrows). HE x400

**Figure 5(c):**
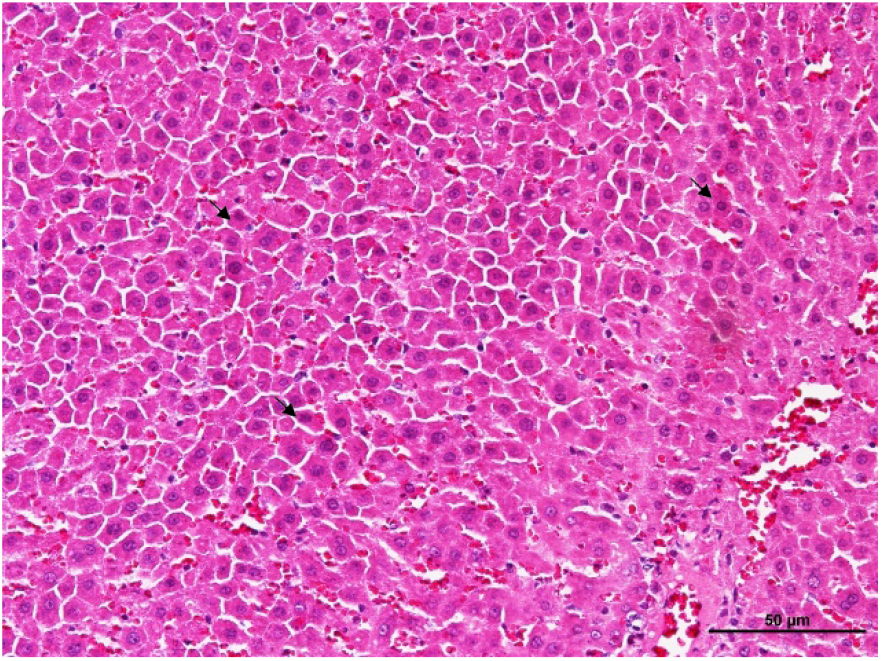
Section of liver of a rat from group CD showing hepatocytes with eosinophilic cytoplasm and pyknotic nuclei (arrows). HE x200

**Figure 5(d):**
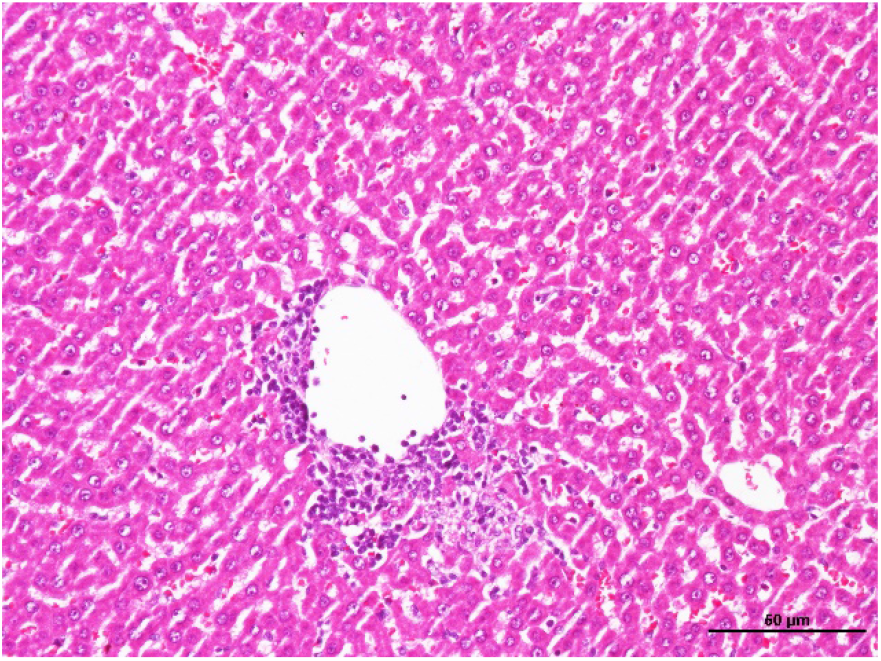
Section of liver of a rat from group CD showing focal hepatitis around a vein. HE x200

**Figure 5(e):**
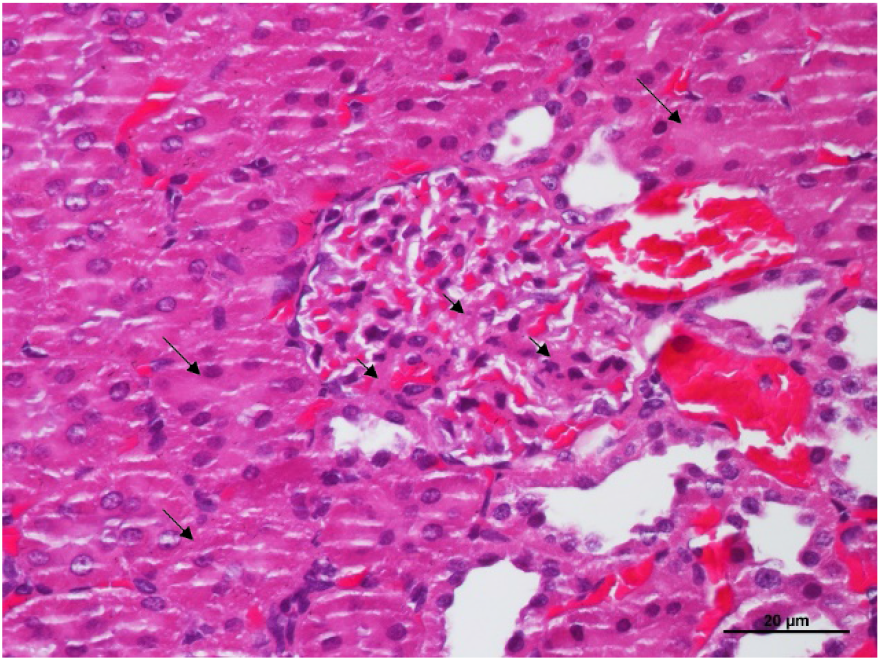
Section of a kidney of rat from group CD showing congested capillaries and thickened mesangium (arrowheads) and basement membrane. Note the epithelium of the convoluted tubules are eosinophilic and swollen, blocking the lumen (arrows) HE x400

**Figure 5(f):**
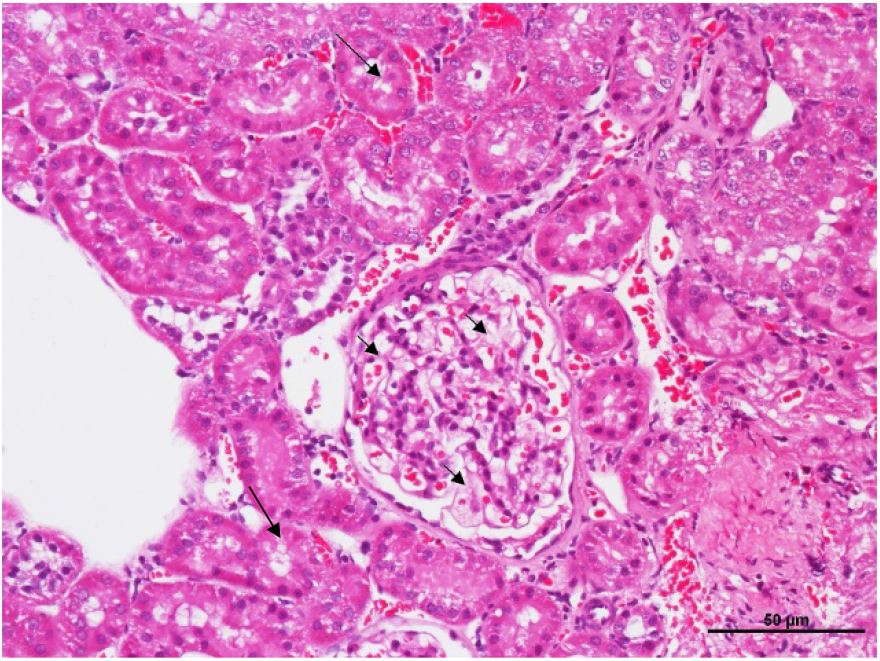
Section of a kidney of rat from group RS2 showing fairly normal capillaries, mesangium (arrowheads) and basement membrane. Note the epithelium of the convoluted tubules are less swollen while the lumens are visible (arrows) HE x200

#### The liver

The liver sections of all groups showed various degree of congestion and thrombosis in the blood vessels, particularly the minor blood vessels. All rats of CD group showed moderate congestion of blood vessels, especially the small vessels, some with thrombus. Many focal areas with hepatocytes that appeared with eosinophilic cytoplasm and pyknotic nuclei **(Figure 5c)**. There were numerous foci of hepatitis with infiltration of mononuclear cells, particularly surrounding some small blood vessels **(Figure 5d)**. Approximately 40% of the hepatocytes showed these changes. Similarly, group HM showed moderate congestion of the small blood vessels, some with thrombus. There were few small foci of hepatitis around small blood vessels with approximately 20% of the hepatocytes, particularly near the hepatitis area showing eosinophilic cytoplasm and pyknotic nucleus. On the other hand, groups sago RS2 and RS4 showed slight congestion of small blood vessels with occasional foci of hepatitis around small blood vessels and bile ducts. Generally, the hepatocytes were fairly normal with occasional small area showing eosinophilic cytoplasm and pyknotic nucelus. Lesions of hepatitis and hepatocyte degeneration were significantly (p<0.05) reduced in groups HM, sago RS2 and sago RS4 **(Table 5)**. Group RS4 showed least extensive hepatocytic degeneration although insignificant (p>0.05). Groups sago RS2 and RS4 showed significantly (p<0.05) less congestion of blood vessels compared to groups CD and HM.

**Table 5:** Lesion scores that indicate the severity of lesion in liver sections

| Group | Lesion score |  |  |
| --- | --- | --- | --- |
|  | Congestion | Hepatitis | Pyknotic |
| Control Diabetic | 1.7±0.5<sup>a</sup> | 2.0±0.9<sup>a</sup> | 2.0±0.9<sup>a</sup> |
| Hi Maize | 2.0±0.0<sup>a</sup> | 1.0±0.0<sup>b</sup> | 1.0±0.0<sup>b</sup> |
| RS2 | 1.0±0.0<sup>b</sup> | 1.0±0.0<sup>b</sup> | 1.0±0.0<sup>b</sup> |
| RS4 | 1.0±0.0<sup>b</sup> | 1.0±0.0<sup>b</sup> | 0.7±0.5<sup>b</sup> |

<sup>a,b</sup>Different superscript represent significant (p<0.05) different when the respective lesion was compared

#### The kidneys

Examinations of the kidneys were focussed on changes in glomerulus, convoluted tubules and blood vessels. In CD group, between 50% and 70% of the glomeruli appeared moderately congested with moderately thickened mesangium and glomerular basement membrane **(Figure 5e)**. Majority of the epithelium that lined the convoluted tubules were swollen thus, closing the lumen of the tubules while the cytoplasm appeared eosinophilic and many nuclei appeared pyknotic. Some of the rats in this groups showed few atrophied glomeruli. Most glomeruli of group HM appeared fairly normal with approximately 30% showing slightly thickened mesangium and basement membrane. Similarly, most convoluted tubules appeared fairly normal with clear lumen and only few tubules with swollen, eosinophilic and pyknotic epithelium. Similarly, groups sago RS2 **(Figure 5f)** and sago RS4 had fairly normal glomeruli with approximately 15% glomeruli with thickened mesangium and basement membrane. Few glomeruli appeared atrophied. Most convoluted tubules appeared normal with 10%-20% with swollen, eosinophilic and pyknotic nuclei. Renal congestion appeared to be significantly (p<0.05) more severe in CD group than other groups **(Table 6)**. Similarly, CD group revealed most severe (p<0.05) glomerular lesions than groups sago RS2 and RS4 while HM showed significantly (p<0.05) least severe glomerular lesions. On the other hand, group sago RS4 showed significantly (p<0.05) least severe lesions in the convoluted tubules compared to groups HM and sago RS2 while CD group showed significantly (<0.05) most severe lesions.

**Table 6:**
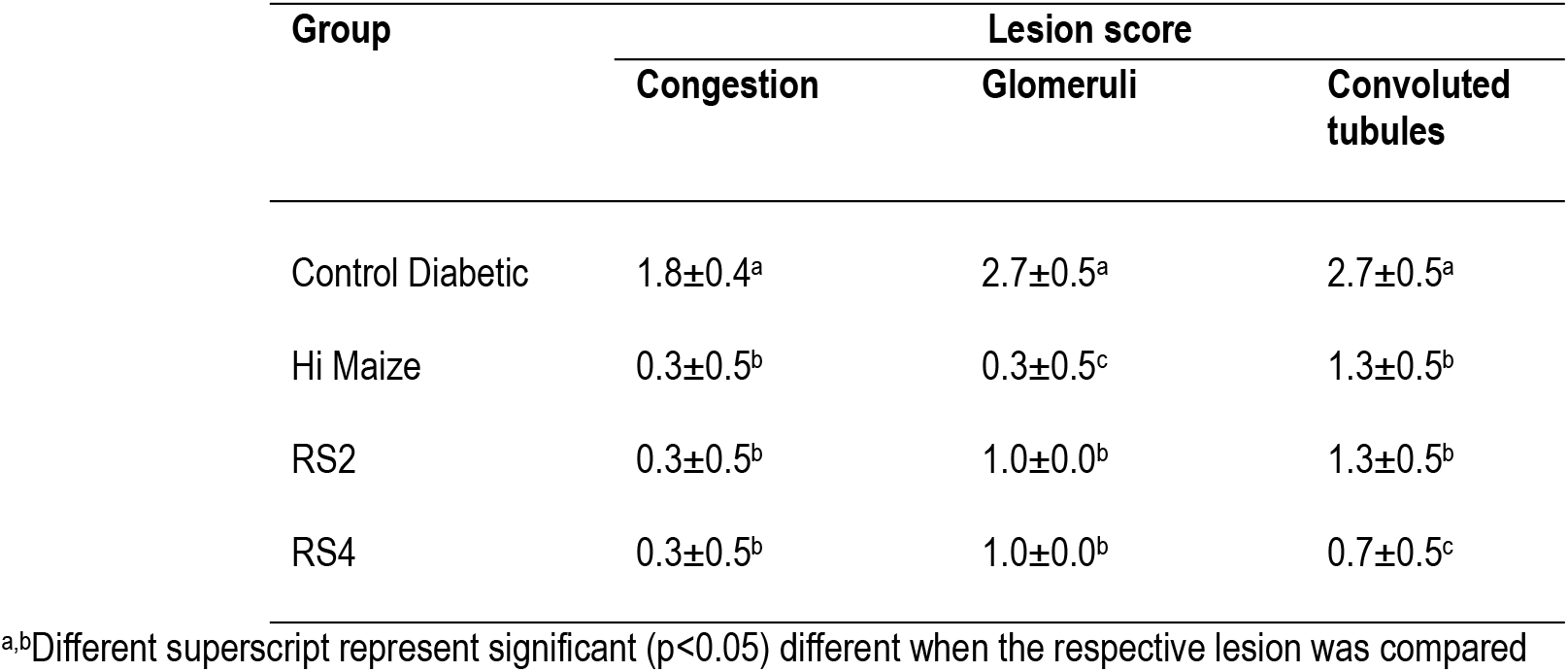
Lesion scores that indicate the severity of lesion in kidney sections

## Discussion

A month treatment with RS in the diabetic Goto kakizaki (GK) rat showed a significant reduction in the area under the glucose curve of the sago RS2 treated group with a similar trend was observed in the sago RS4 and HM groups but not significant. Despite its lower RS content, sago RS2 seemed to be demonstrating better effect in reducing blood glucose as compared to sago RS4. The area under the curve for insulin showed a significant increase in both RS and HM groups which might contribute to the glucose lowering effects in the diabetic GK rat.

Dietary RS resists digestion in the small intestine and is fermented in the large intestine instead, thus preventing blood glucose spike (46). The glucose lowering effect by RS mainly involve the digestion system enzymes by delaying the glucose absorption (47), inhibiting carbohydrate digestion (48) and slowing down gastric emptying (49). Therefore, the mechanisms behind the blood glucose lowering effects by sago RS2 and RS4 in the GK rat might be contributed mainly by α-glucosidase inhibitory activity identified in this study, which act by preventing digestion of starch and glucose release in the intestine (12,13). In addition to this mechanism of action, the DPP(IV) inhibitory effects as shown by Sago RS2 and RS4 may contribute to glucose reduction in the GK rat which act by delaying gastric emptying, stimulating insulin release and reducing glucagon (15).

In principal, inhibition of the enzyme DPP-IV promotes increased activity of incretin hormones including glucagon-like peptide-1 (GLP-1) and glucose-dependent insulinotropic peptide (GIP), thereby regulate glucose by stimulating insulin secretion and reducing postprandial concentrations of glucagon (50). The glucose lowering effect by RS mainly involve the digestive system. However, RS might act partly on insulin via GLP-1 activated by DPP-IV inhibitory activity. Our data showed that improved insulin secretion was observed in both *in vivo* and in *ex vivo* using non diabetic isolated pancreatic in response to sago RS2 and RS4. This is further supported by the increase of GLP-1 hormone in the RS treated group with a significant result shown in the sago RS2. Therefore, the increase of DPP-IV inhibitory activity by RS might triggers GLP-1 level and promotes insulin release. GLP-1 plays an important role for glucose regulation since the hormone improves insulin level and reduce glucagon (44). DPP(IV) inhibitory activity exert its effects via satiety, improved beta-cell production and prevention of beta cells apoptosis (15). However, further study needs to be done to understand the relationship between DPP-IV inhibitory activity and stimulation of insulin release by RS. Glucagon hormone however was not supressed in response to RS as compared to the control group but no significant differences were observed. Therefore, a better understanding on DPP-IV inhibition affecting GLP-1 and glucagon hormones in response to sago RS should be further investigated.

Other hormone, Ghrelin which is an appetite stimulating hormone (51) showed the lowest level in sago RS2 group, which explained the low feed intake. However, in contrast to sago RS2, high level of ghrelin did not increase the overall feed intake in the sago RS4 group. Treatment with sago RS2 and RS4 indicated a significant reduction in body weight compared to control diabetic which could be due to lesser amount of feed and water consumption throughout the study.

In our previous study, we have performed Rat Glucose Metabolism RT^2^ Profiler PCR Array, Qiagen using liver tissues of GK rat treated with RS2 and RS4. Our data showed that multiple genes which are specifically involved in the glucose and glycogen metabolisms were up- and down-regulated indicating the potential role in glucose homeostasis (34). The genes identified might be useful in reducing hepatic glucose output thereby lowering blood glucose level.

Histological examination of pancreas, liver and kidney between different treatments of RS revealed tissue damage characterized by lesion severity. In general, defects in the cells of the Islet of Langerhans are among the major lesions that leads to diabetes (52). In this study, the defect was in the form of necrosis of the cells in which the cells appeared with small, compact, pyknotic nuclei. Control group seemed to show the most severe lesion in the pancreas compared to the treated RS groups although insignificant. Among the consequences of diabetes are the hepatitis and degenerative changes of the hepatocytes in the liver (53) of affected animals and again, control group showed the most severe lesions. One of the most affected organs following diabetes is the kidneys that develop lesions in the glomeruli and convoluted tubules. In this study, group sago RS4 was the only group that showed significantly less severe changes in the convoluted tubules, indicating lesser effect of diabetes on the tubules of kidneys. Treatment with sago RS2 and RS4 seemed to show lesser cells degeneration in the pancreas, liver and kidney tissues examined, therefore, it would be interesting to further identify the markers and underlying mechanisms involved.

## Conclusions

Findings of the study demonstrated that RS2 from Sarawak Sago palm consumption improved glucose tolerance in the diabetic rats indicating a preliminary proof on the health benefit in diabetes management. RS may be potentially developed as a functional ingredient or dietary supplement. The encouraging results justify further evaluation to be carried out in human type 2 diabetes patients, however, the mechanisms of glucose reduction by sago RS need to be further confirmed.

### Statistical analysis

Results were presented as mean ± SEM. The results were evaluated using Kruskal–Wallis test, followed by Dunn’s pairwise test (different groups) and Wilcoxon test (differences within same groups). p-values<0.05 were considered to be statistically significant. All statistical analysis was performed using SPSS 21.0. For DPP-IV inhibitor and α-Glucosidase inhibitory activity, results were evaluated using One-way ANOVA, followed by Tukey’s test for post hoc analysis. For Histology examination, statistical analysis was used to compare the severity of histopathological changes between the different groups before a final conclusion was derived. Student t test was used to compare the results of each treated group.

## Data Availability

The study data used to support the findings are kept by the project team of Downstream Technology Division, CRAUN Research Sdn. Bhd., Sarawak and Endocrine and Metabolic Unit, Institute for Medical Research, Selangor and requests for data will be considered by the data centre.

## Conflict of interest

The authors declare no conflict of interest. CRAUN Research is an agency entrusted by Sarawak State Government to spearhead the Sago Research in Sarawak. This project is a joint-collaboration with the Institute for Medical Research (IMR) of which IMR has assumed full control on the technical conduct of the study.

## Funding

All authors would like to thank the Sarawak State Government for financially funded most of the work involved in this project. The portion of animal study was supported by both CRAUN Research Sdn Bhd, under project number DTD-1.2.2 (59%) and Ministry of Health Research Grant (NMRR-18-3353-45113; 19-013) (41%).

## Acknowledgements

The authors gratefully acknowledge the Sarawak State Government, the Management of CRAUN Research Sdn. Bhd., and the Director General of Health, Malaysia, Director of the Institute for Medical Research, Ministry of Health Malaysia and for permission to publish the finding of this study.

## Authors Contribution

Ezarul Faradianna Lokman, Sal Hazreen Bugam-designed and performed experiments, data analysis, manuscript writing; Nurleyna Yunus, Fazliana Mansor-supervision and idea; Vimala Balasubramaniam, Khairul Mirza Mohamad Rabizah Md Lazim, Aina Shafiza Ibrahim, Awang Zulfikar Rizal Awang Seruji-performed experiments and data analysis

## Notes

### Competing Interest Statement

The authors have declared no competing interest.

